# Collagen modification remodels the sarcoma tumor microenvironment and promotes resistance to immune checkpoint inhibition

**DOI:** 10.1101/2024.06.28.601055

**Authors:** Hehai Pan, Ying Liu, Ashley M. Fuller, Erik F. Williams, Joseph A. Fraietta, T.S. Karin Eisinger

**Affiliations:** Department of Pathology & Laboratory Medicine; Penn Sarcoma Program; Abramson Family Cancer Research Institute; Department of Microbiology; Center for Cellular Immunotherapies; Parker Institute for Cancer Immunotherapy; Perelman School of Medicine; University of Pennsylvania, Philadelphia, PA, USA

## Abstract

Molecular mechanisms underlying immune checkpoint inhibitor (ICI) response heterogeneity in solid tumors, including soft tissue sarcomas (STS), remain poorly understood. Herein, we demonstrate that the collagen-modifying enzyme, procollagen-lysine,2-oxoglutarate 5-dioxygenase 2 (Plod2), which is over-expressed in many tumors relative to normal tissues, promotes immune evasion in undifferentiated pleomorphic sarcoma (UPS), a relatively common and aggressive STS subtype. This finding is consistent with our earlier observation that Plod2 promotes tumor metastasis in UPS, and its enzymatic target, collagen type VI (ColVI), enhances CD8+ T cell dysfunction. We determined that genetic and pharmacologic inhibition of Plod2 with the pan-Plod transcriptional inhibitor minoxidil, reduces UPS growth in an immune competent syngeneic transplant system and enhances the efficacy of anti-Pd1 therapy. These findings suggest that PLOD2 is an actionable cancer target and its modulation could augment immunotherapy responses in patients with UPS, and potentially other sarcomas and carcinomas.

## Introduction

Immunosuppression in the solid tumor microenvironment (TME) arises from mechanisms driven by both the tumor parenchyma and surrounding stroma^1^. One mechanism of particular clinical significance is tumor-induced activation of inhibitory immune checkpoints, mediated by co-receptors expressed on the surfaces of CD8^+^ T cells that attenuate their tumoricidal activity^2, 3^. Accordingly, immune checkpoint inhibitors (ICIs) such as Pembrolizumab (a-programmed cell death protein 1, PD-1) are promising anti-neoplastic agents. However, clinical response rates vary widely (15-60%) among solid tumor patients, with heterogeneous responses reported even within specific cancer types^4^. Thus, a better understanding of ICI response heterogeneity is critical for the development of novel approaches that improve therapeutic efficacy.

Soft tissue sarcoma (STS) is a large, heterogeneous group of solid connective tissue tumors that account for ∼1% of adult cancer cases (up to 350,000 cases worldwide annually)^5, 6^. Undifferentiated pleomorphic sarcoma (UPS) is a relatively common STS subtype that predominantly arises in adult skeletal muscle^7, 8^. Although STS is generally considered non-immunogenic due to its low mutational and neoantigen burden relative to carcinomas, recent clinical trials have revealed that ∼25% of UPS patients exhibit objective clinical responses to a-PD-1 treatment.^9, 10^ These encouraging findings suggest that studies of UPS may provide valuable insights into strategies for enhancing CD8^+^ T cell function and ICI responses in solid tumors.

Defining characteristics of UPS, as well as other sarcomas and high-grade carcinomas, include extensive deposition and aberrant post-translational modification (PTM) of extracellular matrix (ECM) proteins^11-14^. In particular, the solid tumor ECM is rich in molecules belonging to the collagen superfamily, a large, diverse protein family that contains 28 molecular species encoded by over 40 genes^15^. Although the roles of individual collagen proteins in cancer-associated processes are only beginning to be defined - particularly with respect to their impact on solid tumor immune evasion - our recent work demonstrated that collagen type VI (ColVI), a microfibrillar collagen, facilitates tumor progression by inhibiting T cell migration, infiltration, and function^16, 17^. Specifically, ColVI promotes immune evasion and hinders ICI efficacy in UPS by inactivating CD8^+^ T cells and remodeling collagen type I (ColI) in the TME^16^. These findings implicate ColVI ablation, in combination with immunotherapy, as a promising approach to mitigate immune evasion in UPS, and potentially other solid tumors. As ColVI molecules in the UPS TME are unlikely to be directly “druggable”, investigation of ColVI-interacting proteins may reveal potential ColVI-targeting strategies. The lysine hydroxylase procollagen-lysine,2-oxoglutarate 5-dioxygenase 2 (PLOD2) is a collagen-modifying enzyme required for normal collagen production across many tissue types; germline *PLOD2* mutations result in Bruck syndrome, a rare, autosomal-recessive disease in which patients present with osteoporosis, scoliosis, and joint contractures due to under-hydroxylated COLI molecules^18, 19^. In malignant contexts, we and others have established that *PLOD2* overexpression promotes metastasis and reduces long-term survival in the setting of UPS and various epithelial tumors (e.g., bladder, liver, renal, and lung), in part through aberrant hydroxylation of ColVI in the TME^20-28^. However, because these prior *in vivo* studies were predominantly carried out in immunodeficient xenograft models, the role of cancer cell-intrinsic Plod2 in primary tumor immune evasion remains underexplored. Therefore, herein, we queried the role of UPS cell-intrinsic Plod2 in immunosuppression and adaptive immune cell function in the UPS TME. Our findings suggest targeting collagen modification as a novel approach to potentiate ICI efficacy in UPS, and potentially other solid tumors.

## Methods

### Cell lines

Human STS-109 cells were derived from a pre-treatment human UPS tumor by Rebecca Gladdy, M.D. (University of Toronto) and cultured in DMEM with 20% (vol/vol) FBS, 1% L-glutamine, and 1% penicillin/streptomycin. HEK-293T cells were purchased from ATCC (Manassas, VA, USA) and cultured in DMEM with 10% (vol/vol) FBS, 1% L-glutamine, and 1% penicillin/streptomycin. Murine sarcoma SKPY42.1 cells on a C57BL/6 background (a gift from Sandra Ryeom, PhD, Columbia University) were cultured in DMEM with 10% (vol/vol) FBS, 1% L-glutamine, and 1% penicillin/streptomycin. All cell lines were cultured at 37°C in 5% CO_2_ and confirmed to be negative for mycoplasma contamination.

### Murine models

All animal experiments were performed in accordance with NIH guidelines and approved by the University of Pennsylvania Institutional Animal Care and Use Committee. We generated genetically engineered mouse model (GEMM) tumors by injecting a calcium phosphate precipitate of adenovirus expressing Cre recombinase (University of Iowa) into the right gastrocnemius muscle of 3-month-old LSL-*Kras*^G12D+^; *Trp53*^fl/fl^ (KP) mice as previously described^29^. For syngeneic transplant studies, 1 million SKPY42.1 cells were resuspended in 100 ul PBS and implanted subcutaneously into syngeneic 5-6-week-old C57BL/6 mice (The Jackson Laboratory; strain code 000664).

### In vivo drug treatment, tumor growth measurements, and survival analyses

For minoxidil-alone studies, minoxidil (Sigma-Aldrich, M4145) or vehicle control (PBS) was administered intraperitoneally (I.P.) daily at 10 mg/kg or 30 mg/kg, beginning 12 days after initial implantation of UPS cells. In immune checkpoint studies, 200 μg of anti-Pd1 monoclonal blocking antibody (BE0146, BioXCell) or isotype control antibody (BE0089, BioXCell) was administered I.P. every three days once tumors became palpable. Minoxidil (or vehicle control) was administered at 30 mg/kg in all studies evaluating efficacy in combination with anti-Pd1 antibodies. In all studies, tumors were measured every 2-3 days using calipers, and volumes were calculated using the formula (ab^2^)π/6, where a and b indicate the longest and shortest dimensions, respectively. Body weights were recorded every 2-3 days, and tumor volumes of 2000 mm^3^ were used as endpoints for survival analysis.

### Two photon second-harmonic generation (SHG) imaging of tumor collagen fibers

Organization of collagen fibers in formalin-fixed, paraffin-embedded murine tumor sections were visualized with second harmonic generation (SHG) imaging using a Leica TCS SP8 MP 2-photon microscope (Leica Microsystems) as previously described^16^. The unstained tissue slides were maintained in water, and image stacks (6.99 μm) were acquired using a 25X 1.0NA water immersion objective with 4x zoom. Collagen fiber width and orientation distribution from SHG image stacks (maximum-intensity projections) were quantified using CT-FIRE as described previously^16, 30^.

### RNA isolation and RT-qPCR

Total RNA was isolated from cells using the RNeasy Mini Kit (Qiagen, Cat #74104) and tissue using TRIzol (Thermo Scientific, #15596018). Reverse transcription of mRNA was performed using the High-Capacity RNA-to-cDNA Kit (Thermo Scientific, #4387406). RT-qPCR was performed using a ViiA7 real-time PCR system (Applied Biosystems). All probes were TaqMan “Best coverage” probes (Life Technologies). *HPRT1* was used as a normalization control.

### Lentiviral Transduction

Glycerol stocks for human *PLOD2*-targeting (TRCN0000064809, TRCN0000064808), murine *Plod2*-targeting (TRCN0000076409, TRCN0000076411), and scrambled shRNAs were obtained from Sigma Aldrich. shRNA plasmids were packaged using the third-generation lenti-vector system (pMDLg/pRRE, pRSV-Rev, and pMD2.G/VSVG) and expressed in HEK-293T cells. Supernatant was collected at 24 and 48 hrs after transfection and subsequently concentrated using polyethylene glycol-8000. Virus transduction of target sarcoma cells was performed in the presence of 8.0 μg/mL polybrene (Sigma-Aldrich, #H9268) and puromycin selection (3 μg/mL) was performed after 48 hours. RT-qPCR was performed to confirm knockdown efficiency of target genes at day 5 after initial viral transduction.

### Human CAR-T cell production and xCELLigence real-time cytotoxicity assay

The xCELLigence real-time cytotoxicity assay was performed as previously described.^16^ In brief, STS109 cells (target cells) expressing a control or one of multiple *PLOD2*-targeting shRNAs were harvested and seeded in triplicate into 96-well polyethylene terephthalate plates (E-Plate VIEW 96 PET, Agilent) at 20,000 cells/per well. STS-109 cells were allowed to adhere to the bottoms of wells for 24 hours (37°C, 5% CO_2_ conditions) after which effector cells (CART-TnMUC1 cells or donor-matched un-transduced T cells [NTD cells]) were added to wells at effector:target cell ratios of 10:1, 5:1, 1:1, and 0:1. Cytotoxic activity of the T lymphocytes was determined via continuous acquisition of impedance data over 6 days. Raw impedance data was analyzed using 762 RTCA 2.1.0 software as described in^16^.

### Human samples

De-identified human sarcoma and non-malignant muscle samples were obtained from surgically resected tumors from patients undergoing therapeutic surgical resection in accordance with protocols approved by the Institutional Review Board at the University of Pennsylvania.

### Computational analyses of human samples

Associations between *PLOD2* expression levels and human patient survival in The Cancer Genome Atlas-Sarcoma (TCGA-SARC) dataset were queried with the cBioPortal web server (https://www.cbioportal.org/). The publicly available Affymetrix Human Genome U133A microarray dataset reported in Detwiller et al.^31^; NCBI Gene Expression Omnibus accession number GSE2719) was used to compare *PLOD2* gene expression levels in sarcoma and normal connective tissue. Data pre-processing was carried out in RStudio as follows: Data were quantile normalized, and duplicate probes corresponding to the same ENTREZ gene ID were collapsed by averaging. The resulting gene expression values were log-transformed with a pseudocount of 1 and median centered across samples.

### Statistical analysis

Statistical analysis was performed using GraphPad Prism (version 10). Data are shown as mean ± SEM or SD as indicated in the figure legends. Student t-tests (unpaired two-tailed) were performed to determine whether the difference between two means was statistically significant. ANOVA was used for such assays with three or more groups. 2-way repeated-measures ANOVA, mixed-effects models, or non-linear regression models (exponential fit) were used for analyses of *in vivo* tumor growth curves.

## Results

### Plod2 promotes UPS primary tumor growth in an immunocompetent setting

Previous reports investigating the role of cancer cell-intrinsic Plod2 in primary tumor growth have primarily relied on xenograft studies in nude mice. These animals lack mature adaptive immune cells (T cells and B cells) but generally retain innate immunity (e.g., myeloid-lineage cells). Therefore, to understand the role of UPS cell-intrinsic Plod2 in immune evasion, we leveraged the *Kras*^*G12D/+*^; *Trp53*^*fl/fl*^ (KP) syngeneic transplant system previously introduced in^16^. In this system, sarcoma cells derived from the gold-standard KP genetically engineered mouse model (GEMM) of UPS^29^ on a pure C57BL/6 background (herein referred to as “KP cells”) are implanted subcutaneously or orthotopically (intramuscularly) into syngeneic, immunocompetent C57BL/6 mice.^16^ To this end, we transduced KP cells with a control (shScr) or one of multiple *Plod2-*targeting (shPlod2) shRNAs, injected them subcutaneously into recipient animals, and tracked tumor growth (**Fig. 1A-C**). Remarkably, unlike in previous work where UPS cell-intrinsic *Plod2* depletion had no impact on primary tumor progression in immunodeficient nude mice^20^, *Plod2*-deficient UPS tumors were substantially smaller than *Plod2-*sufficient controls in an immunocompetent setting. To validate this genetic observation with a pharmacologic approach, we treated syngeneic KP UPS tumor-bearing mice with minoxidil, a non-specific transcriptional inhibitor of Plod2 and its homologs Plod1 and Plod3^32^; **Fig. 1D**). Minoxidil induced modest dose-dependent reductions in UPS tumor growth, confirming that Plod2 depletion can suppress primary tumor progression in the context of an intact immune system. Together, these results suggest that the presence of adaptive immune cells is required for Plod2-dependent UPS tumor growth, and that cancer cells expressing high levels of Plod2 may promote immunosuppression.

**Figure 1:**
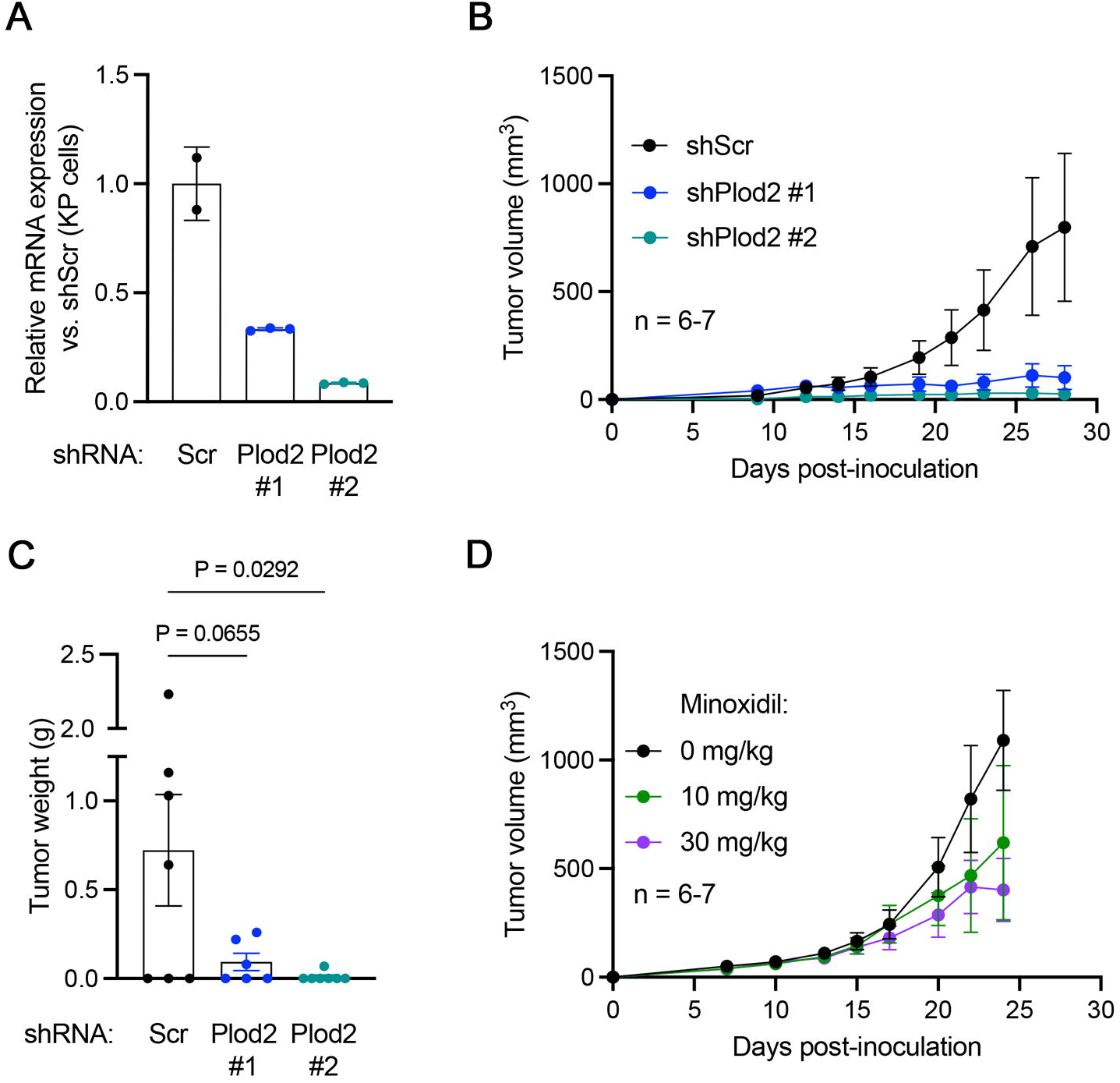
UPS cell-intrinsic Plod2 promotes primary tumor growth in an immunocompetent setting. **A**. Validation of *Plod2* expression in KP UPS cells (SKPY42.1 cell line) prior to *in vivo* implantation. Mean ± SD. Technical replicates; no statistics are shown. **B**. Tumor growth curves from subcutaneous syngeneic transplant of 1 × 10^6^ KP cells from **A** in C57BL/6 mice. Not significant by two-way repeated measures ANOVA with Dunnett’s post-hoc test (vs. shScr). **C**. Weights of excised tumors from **B**. One-way ANOVA with Dunnett’s post-hoc test (vs. shScr). **D**. Tumor growth curves of subcutaneous syngeneic KP tumor-bearing mice treated daily (i.p.) with 10 mg/kg or 30 mg/kg minoxidil, beginning 12 days after UPS cell implantation. Not significant (mixed-effects model with Dunnett’s post-hoc test vs. shScr). For **B-D**, error bars indicate mean ± SEM.

### Plod2 inhibition potentiates immune checkpoint inhibitor efficacy in UPS

To further investigate the relationship between UPS cell-intrinsic PLOD2 and suppression of adaptive immunity, we explored the effects of PLOD2^+^ UPS cells on T cell function. We focused on T cells in these assays, rather than B cells, because T cells are critical players in endogenous anti-tumor immune responses and key targets of immunotherapy strategies^33^. First, we leveraged Tn-MUC1 chimeric antigen receptor T cells (Tn-MUC1 CAR T cells), which target the Tn isoform of mucin 1, a cancer neoantigen^34^. We co-cultured these cells with Tn-MUC1^+^ primary human STS-109 UPS cells^16^ expressing a control or one of multiple *PLOD2-*targeting shRNAs at multiple effector:target ratios, and tracked longitudinal UPS cell lysis (**Figure 2A-B**). This experiment revealed that T cell cytolysis was enhanced in the presence of *PLOD2-*deficient UPS cells, confirming that UPS cells expressing high levels of *PLOD2* suppress T cell function. To explore this relationship *in vivo*, we treated mice bearing syngeneic subcutaneous KP tumors with minoxidil, alone or in combination with ɑ-Pd1 checkpoint therapy. We hypothesized that minoxidil would augment the efficacy of immune checkpoint blockade due to enhanced T cell function in the setting of attenuated UPS cell-intrinsic Plod2 expression. Consistent with this hypothesis, combination treatment with minoxidil and ɑ-Pd1 significantly impaired primary UPS tumor progression compared to treatment with either monotherapy alone (**Figure 2C**). We confirmed this observation in the gold-standard autochthonous KP GEMM, in which mice receiving combination therapy (minoxidil + ɑ-Pd1) exhibited increased survival times (time-to-maximum tumor volume) compared to animals receiving single-agent treatments (**Figure 2D**). Thus, we conclude that pharmacologic Plod2 suppression potentiates the efficacy of ɑ-Pd1 checkpoint therapy in UPS.

**Figure 2:**
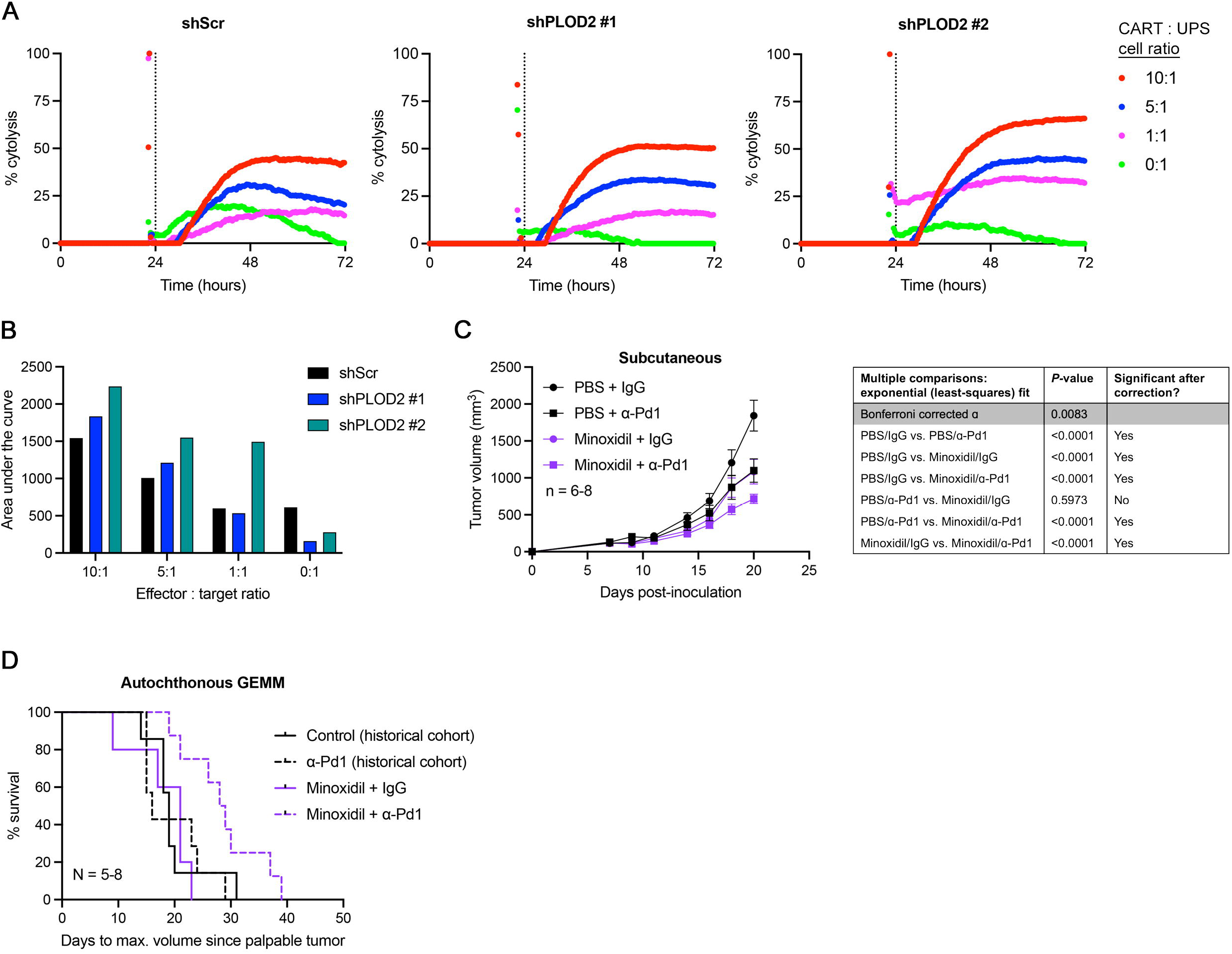
Pharmacologic inhibition of Plod2 potentiates the efficacy of ɑ-Pd1 checkpoint therapy in UPS. **A**. Representative longitudinal cytolysis curves of shScr or shPLOD2 human STS-109 UPS cells co-cultured with CART-TnMUC1 cells from 1 independent human donor. Measurements indicate percent target (UPS) cell cytolysis. **B**. Quantification of data from **A**. Statistics are not shown because n < 3 (n = 2). **C**. Growth curves of subcutaneous (flank) syngeneic tumors in C57BL/6 mice treated with minoxidil (or vehicle control), alone or in combination with ɑ-Pd1 (or isotype control) antibodies. Growth curves were fit with non-linear regression (exponential fit) models; pairwise curve fitting comparisons (extra sum-of-squares F test) are shown in the table at right. **D**. Kaplan-Meier survival curves of KP mice treated with minoxidil, alone or in combination with α-Pd1 checkpoint therapy. Statistics are not shown because data from the control and α-Pd1-alone groups are from a historical cohort first reported in.^16^

### PLOD2 inhibition alters fibrillar collagen architecture in UPS

Previous reports have shown that fibrillar collagen organization is a critical determinant of CD8^+^ T cell activation/function in the UPS TME^16^, and that the lysyl hydroxylase activity of Plod2 is critical for maintaining the structure of mature collagen molecules^21^. Therefore, we explored the effects of Plod2 inhibition on collagen fiber organization in the UPS ECM. To this end, we treated mice bearing syngeneic subcutaneous KP tumors with minoxidil or vehicle control and analyzed the architecture of fibrillar collagen molecules in excised tumor sections using multiphoton second-harmonic generation (SHG) imaging. We observed that tumors from minoxidil-treated animals exhibited significantly thinner fibers than those from the control group (**Fig. 3A-B**). Moreover, by measuring the fiber angle relative to the channel direction (x-axis) and plotting the angle frequency against its distribution, we determined that a substantial portion of collagen fibers in control tumors were aligned with each other in a parallel orientation (**Fig. 3C-E**). In contrast, fibers in minoxidil-treated tumors appeared to have a less uniform orientation. Thus, we conclude that Plod2 inhibition remodels fibrillar collagen molecules in the UPS TME, with possible implications for CD8^+^ T cell function.

**Figure 3:**
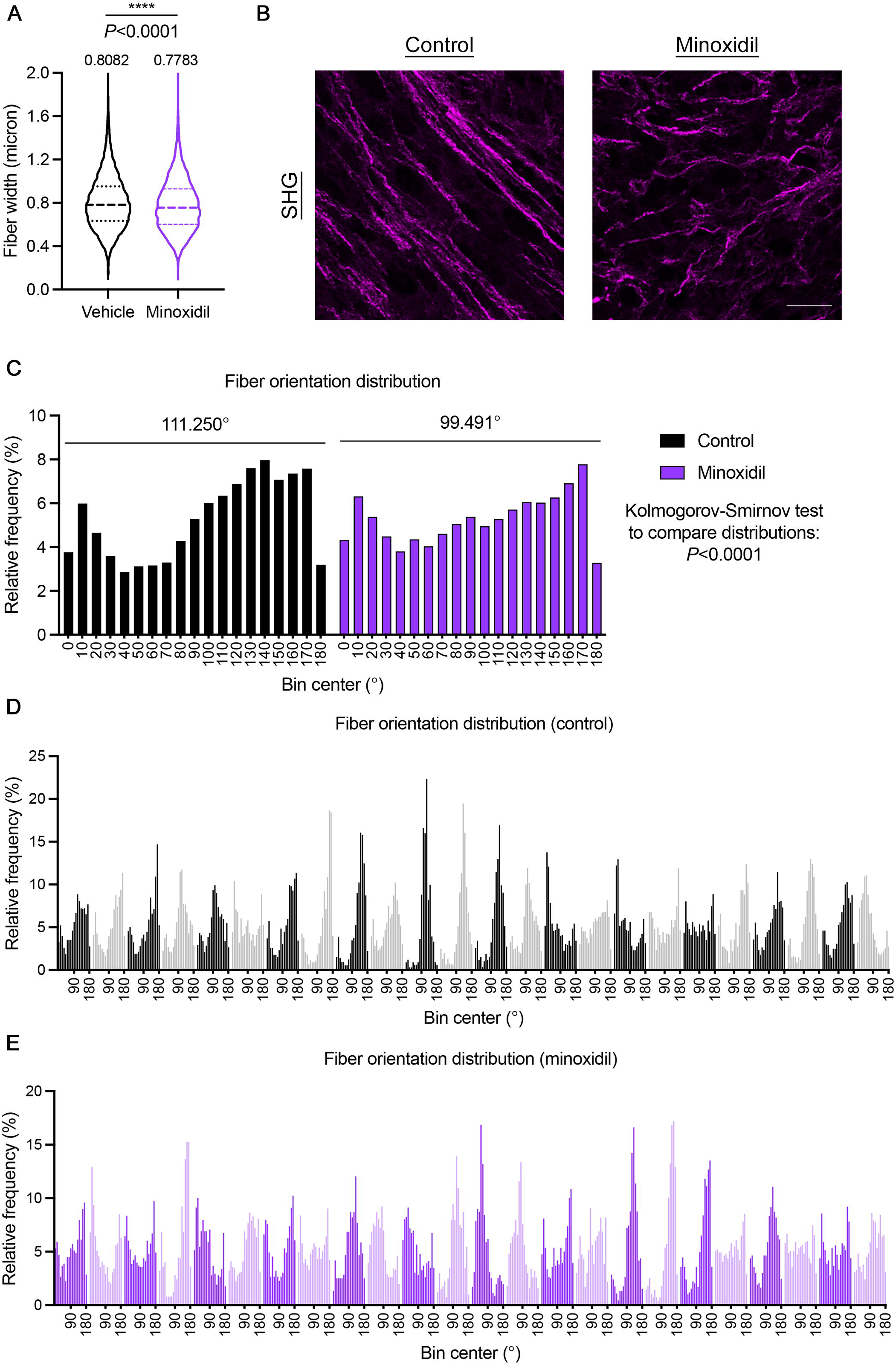
Pharmacologic *PLOD2* inhibition alters fibrillar collagen architecture in UPS. **A**. Violin plots depicting CT-FIRE analysis of fibrillar collagen width from tumor sections from mice bearing subcutaneous syngeneic KP tumors treated with minoxidil or control (second harmonic generation [SHG] imaging). Numbers above violin plots indicate means. Thick and thin dotted lines within each plot denote medians and quartiles 1 and 3, respectively. Two-tailed unpaired t-test. **B**. Representative SHG images of tumors from **A**. Brightness and contrast have been adjusted for presentation purposes. **C**. Frequency distribution histograms of fibrillar collagen fiber orientation in tumors from **A-B** (all microscopy fields combined). Kolmogorov-Smirnov test to compare distributions. **D-E**. Frequency distribution histograms of fibrillar collagen fiber orientation in individual microscopy fields of control (**D**) and minoxidil-treated (**E**) tumors from **C**. Alternating light and dark colors in each plot are meant to enable visual separation of individual fields. For **A-E**, fiber width and orientation were plotted from 6 independent fields across 4 tumors per condition (total n = 24 images/condition).

### PLOD2 is highly expressed and associated with a poor prognosis in STS

To understand the potential clinical utility of PLOD2 inhibitors for UPS, we characterized PLOD2 gene expression patterns in UPS and normal connective tissue specimens. We also included samples from other sarcoma subtypes in this analysis to explore the potential applicability of this approach to a broader swathe of patients with mesenchymal tumors. Using data from multiple human patient cohorts, including the Detwiller et al. dataset^31^ and surgical specimens from the Hospital of the University of Pennsylvania (HUP), we observed that *PLOD2* gene expression levels were generally upregulated in UPS relative to normal connective tissue samples (**Fig. 4A-B**). Conversely, *PLOD2* levels in other sarcoma subtypes such as myxofibrosarcoma (MFS), liposarcoma (LPS), and synovial sarcoma (SS) were more heterogeneous. Consistent with these observations, high levels of *PLOD2* expression were associated with significantly reduced disease-free, disease-specific, and overall survival among UPS patients in The Cancer Genome Atlas-Sarcoma (TCGA-SARC) dataset (**Fig. 4C**). Similar, albeit attenuated, relationships were observed among all TCGA-SARC patients (**Fig. 4D**). Thus, modulation of *PLOD2* expression and/or activity may improve clinical outcomes in UPS patients, and potentially those with other sarcoma subtypes.

**Figure 4:**
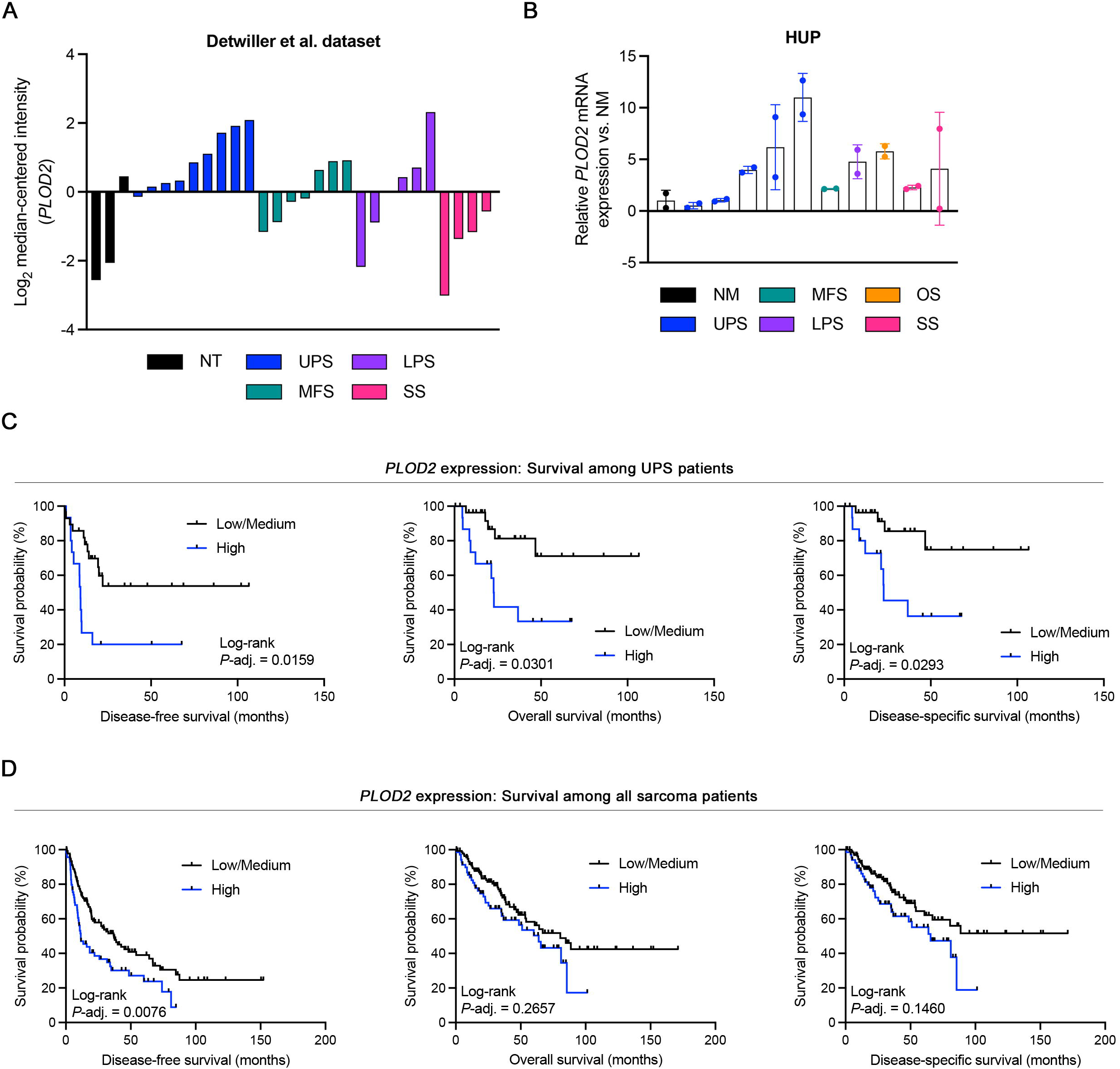
PLOD2 is highly expressed and associated with a poor prognosis in STS. **A**. *PLOD2* gene expression levels in sarcoma and normal connective tissue specimens from the Detwiller et al. data set.^31^ MFS: myxofibrosarcoma; LPS: liposarcoma; SS: synovial sarcoma. **B**. qRT-PCR analysis of *PLOD2* expression levels in human sarcoma and normal skeletal muscle tissue specimens (Hospital of the University of Pennsylvania [HUP]). **C**. Kaplan-Meier survival curves of UPS patients in TCGA-SARC stratified by intratumoral *PLOD2* gene expression levels. **D**. Kaplan-Meier survival curves of all patients in TCGA-SARC stratified by intratumoral *PLOD2* gene expression levels.

## Discussion

Over the last 20 years, studies in multiple cancer types have shown that secretion of matrix proteins into the primary tumor milieu impacts tumor progression^35, 36^. Specifically, collagen fibers can physically associate with cancer cells and promote their migration and invasion^36^. Collagens are also capable of signaling to other cells in the microenvironment including immune cells and platelets^16, 37-39^. In many carcinomas, recruited stromal cells, including activated fibroblasts, secrete ECM proteins^40^. More aggressive epithelial cancer cells, particularly those that have undergone epithelial to mesenchymal transition (EMT), often can secrete ECM as well.^41^ However, given the mesenchymal origins of sarcomas, these cells are intrinsically migratory and invasive and secrete large amounts of matrix, promoting metastasis^20^.

Critically, matrix protein function in both normal and tumor tissues is influenced primarily by significant PTMs.^19, 20, 36, 42-44^ These PTMs dramatically alter collagen structure, organization, and signaling to various cell types. In particular, the hypoxia-inducible collagen-modifying enzyme PLOD2 is a collagen lysyl hydroxylase required for collagen production in normal tissues. However, aberrant *PLOD2* overexpression in solid tumors such as UPS, as well as bladder, breast, liver, and other carcinomas, promotes cancer cell metastasis and is associated with reduced long-term patient survival^24, 45, 46^. Genetic inhibition of *Plod2* and treatment with the pan-Plod inhibitor minoxidil dramatically inhibits metastasis in immunodeficient models.^20^ Mechanistically, excessive collagen lysyl hydroxylation due to hypoxic induction of PLOD2 results in secretion of immature collagen aggregates, which cancer cells can utilize to facilitate their migration toward blood vessels and entrance (intravasation) into the vasculature^20^. Aberrant *Plod2* expression also promotes secretion of lysyl-hydroxylated ColVI into the vasculature, promoting lung endothelial barrier dysfunction and metastasis^21^.

However, the effects of UPS cell-derived Plod2 on collagen signaling to immune cells have remained unclear. Recently, several groups have identified mechanisms by which matrix organization and turnover impact immune evasion in the TME^16, 47-49^. These findings led us to query the impact of aberrant PLOD2 expression in primary UPS tumors on the activation status of CD8+ T cells. Here, we report that *Plod2* genetic depletion or pan-Plod pharmacologic inhibition in syngeneic UPS allografts abolished primary tumor growth. Importantly, Plod2-dependent tumor growth is unique to immune-competent systems. Specifically, Plod2-deficient UPS xenografts implanted in B and T cell-deficient nude mice grow at the same rate as controls^20^, suggesting that Plod2 suppresses the function of the adaptive immune system in the TME of primary UPS tumors. Consistent with these observations, high *PLOD2* expression in human UPS is associated with poor survival. These critical findings suggest that PLOD2 inhibition may augment ICI therapy in UPS patients.

Recent clinical trials have demonstrated that the overall response rate of UPS patients to ICI (Pembrolizumab) treatment is only ∼25%^10, 50^. Development of novel strategies to improve ICI efficacy would be transformative for patients with UPS, and potentially other solid tumors. However, molecular mechanisms underlying ICI response heterogeneity in solid tumors, including UPS, remain poorly understood. Herein, we have demonstrated that sarcoma cell-intrinsic *PLOD2* suppresses T cell function using a TnMUC1-CART system. We also tested the effect of pan-PLOD inhibition combined with ICI in both syngeneic and GEMM models of UPS, and found that combination treatment was substantially more effective than either treatment alone. These findings are consistent with the idea that PLOD2 promotes UPS tumor growth by enhancing T cell dysfunction and immune evasion. Our data also implicate PLOD2-mediated alterations in collagen organization as a potential mechanism underlying PLOD2-driven CD8+ T cell inactivation. Therefore, targeting collagen-modifying enzymes such as PLOD2 represents a potential actionable intervention for augmenting immunotherapy responses in UPS patients. In addition to pursuing PLOD2 inhibitors, which are currently in development^51^, there is also a need for an appropriate biomarker to identify appropriate patient cohorts for PLOD2-targeting therapies. Some biomarker possibilities include levels of PLOD2 expression in primary tumor biopsies by IHC, levels of circulating collagens such as the oncogenic PLOD2 substrate COLVI^16, 21^, or novel approaches to detect excess lysyl hydroxylation of collagen in tissue or blood.

Ultimately, the studies described here support the development of matrix-specific and novel immunotherapy strategies for patients with UPS, and potentially other sarcomas and carcinomas.

## Acknowledgements

We would like to thank James Hayden and Frederick Keeney of the Wistar Institute Imaging Facility for their assistance with multiphoton microscopy and analysis. This work was funded by The University of Pennsylvania Abramson Cancer Center, The Penn Sarcoma Program, Steps to Cure Sarcoma, DoD IDA award RA200237, and NIH/NCI R01CA229688.

## Author contributions

Conceptualization: TSKEM, YL, HP

Methodology: AMF, HP, YL, EFW

Validation: AMF, HP, YL,

Formal Analysis: AMF, HP, YL, EFW

Investigation: HP, AMF, YL, EFW

Data Curation: AMF, YL

Provision of resources: JAF, TSKEM

Writing-original draft preparation: AMF, HP, YL, TSKEM

Writing-review and editing: AMF, HP, YL, TSKEM

Visualization: AMF, YL, TSKEM

Supervision: JAF, TSKEM

Project administration: TSKEM

Funding acquisition: TSKEM

